# Impact of temperature on patient-derived dengue virus breakthrough infections in wMel-infected Aedes aegypti

**DOI:** 10.64898/2026.04.13.718183

**Authors:** Daniela da Silva Gonçalves, Vi Tran Thuy, Robson Kriiger Loterio, Nhu Vu Tuyet, Trang Huynh Thi Xuan, Giang Nguyen Thi, Van Huynh Thi Thuy, Huynh LeDuyen, Dui Le Thi, Long Thi Vo, Huynh Le Anh Huy, Nguyen Thi Van Thuy, Phong Thanh Nguyen, Sophie Yacoub, Katherine L. Anders, Heather Flores, Cameron P. Simmons, Johanna E. Fraser

**Affiliations:** Oxford University Clinical Research Unit, Ho Chi Minh City 710400, Vietnam; Department of Microbiology, Monash Biomedicine Discovery Institute, Monash University, Clayton, VIC 3800, Australia; Life Sciences Discipline, Health Security and Pandemic Preparedness Program, Burnet Institute, Melbourne, Victoria, Australia Burnet Institute, Melbourne, VIC 3004, Australia; World Mosquito Program, Monash University, Clayton, VIC 3800, Australia; School of Public Health and Preventive Medicine, Monash University, Melbourne, VIC 3004, Australia; School of Biological Sciences, Monash University, Melbourne, Victoria, Australia

## Abstract

The *w*Mel strain of the insect endosymbiont *Wolbachia* reduces the potential for *Aedes aegypti* to transmit mosquito-borne viruses such as dengue (DENV). Field trials that have introgressed *w*Mel into *Ae. aegypti* populations have shown this approach significantly reduces dengue incidence. In a laboratory setting some *w*Mel-*Ae. aegypti* develop infectious saliva following a viremic blood meal. Additionally, studies have demonstrated that exposing *w*Mel-*Ae. aegypti* to heat treatment, particularly during the larval stage, reduces *w*Mel density in key tissues such as the ovaries, midgut and salivary glands. Here we build on these studies, using viremic blood collected from 13 dengue inpatients at the Hospital for Tropical Diseases in Ho Chi Minh City (Viet Nam), to assess how temperature affects the protection afforded to *Ae. aegypti* by *w*Mel.

We found that, compared to *w*Mel-*Ae. aegypti* reared at 28 ± 4°C, those reared at 31 ± 4°C developed infectious saliva more frequently, but the risk of this occurring was still reduced compared to WT mosquitoes reared at the same temperature. Heat treatment reduced the density of *w*Mel in all tissues tested, decreased the magnitude of *w*Mel’s protection against DENV replication in the head/thorax, and significantly increased the amount of DENV replication in *w*Mel-*Ae. aegypti*. When comparing cohorts of *w*Mel-*Ae. aegypti* that did or did not develop infectious saliva, DENV levels in the head/thorax were associated with increased odds of mosquitoes developing infectious saliva, but *w*Mel density was not.

Overall, these findings show that elevated rearing temperatures increase the risk of patient-derived DENV breakthrough infections in *w*Mel-*Ae. aegypti*, potentially due to increased DENV replication in these mosquitoes. This limitation suggests it would be prudent to increase surveillance in regions using *w*Mel for dengue control when daily mean temperatures remain above 30°C for multiday periods.

**Author Summary:** The mosquito species *Ae. aegypti* can be infected with the bacterium *Wolbachia* (*w*Mel strain), reducing its capacity to transmit viruses like dengue (DENV). *Wolbachia* is now being used as a biocontrol tool to reduce the burden of dengue in communities. However, some mosquitoes with *Wolbachia* can still transmit DENV. Here we utilised a natural infection model using dengue patient-derived blood to examine how temperature may increase the risk of DENV transmission occurrence in mosquitoes with *w*Mel. Mosquitoes with *w*Mel were more likely to transmit virus when reared at an average temperature of 31°C compared to those reared at an average temperature of 28°C but these mosquitoes still had a lower risk of developing infectious saliva compared to their *w*Mel-free counterparts. Higher temperatures reduced the amount of *w*Mel in mosquito tissues and increased the amount of DENV replicating in the head/thorax. Increasing levels of DENV RNA in these tissues were found to be associated with increased risk of mosquitoes with *w*Mel developing infectious saliva.

This finding indicates surveillance is warranted in high temperature settings or during heat waves, to monitor for changes in *w*Mel frequency and DENV infection in *Ae. aegypti*.

## Introduction

Mosquito-borne viruses such as dengue (DENV) impose an enormous health and economic burden on populations in tropical and sub-tropical climates around the world (1–3). Dengue incidence continues to rise globally, in part due to urbanisation, international travel, and the increasing geographic spread of the primary mosquito vector, *Aedes aegypti*, which is now found on all inhabited continents (4). Sub-optimal dengue vaccines (5, 6) and the emergence and spread of insecticide resistance in *Ae. aegypti* populations (7) has undermined conventional dengue control efforts.

Over the past decade infection of *Ae. aegypti* with *Wolbachia* has emerged as an effective public health intervention. *Wolbachia* is an endosymbiotic bacterium naturally found in at least 40% of terrestrial arthropods (8), but not *Ae. aegypti* (9). *Wolbachia* strain *w*Mel (originally from *Drosophila melanogaster*) has been stably transinfected into *Ae. aegypti* where it restricts replication of positive-sense RNA viruses like DENV (10). *Ae. aegypti* infected with *w*Mel have a reduced risk of transmitting all four serotypes of DENV (DENV-1-4) (10–13).

To implement *w*Mel as a biocontrol tool, *w*Mel-infected *Ae. aegypti* are periodically released into selected dengue-prone regions. *w*Mel spreads through the local *Ae. aegypti* population due to its maternal transmission and ability to reproductively manipulate its host (14–17). Multiple field trials have measured significantly reduced dengue incidence in communities following the establishment of *w*Mel-infected *Ae. aegypti* (18–21). While studies to date suggest *w*Mel’s genome and antiviral activity are quite stable (22–24), there remains a hypothetical risk that DENV will evolve and overcome its antiviral effects (25).

*Ae. aegypti* can become infected with DENV when the mosquito ingests a blood meal from a viremic person. Virus escapes the midgut, passing into the haemocoel where it can infect and replicate in tissues and organs throughout the mosquito. By 7-10 days post blood meal, virus may have disseminated to mosquito salivary glands, allowing the virus to be transmitted onwards as the mosquito probes and salivates whilst blood feeding (26, 27).

In laboratory experiments, when *w*Mel-*Ae. aegypti* are fed blood spiked with cell culture-derived DENV, a proportion of mosquitoes still establish infection in the midgut, but the probability of midgut infection and the amount of virus that disseminates to distal tissues is substantially reduced compared to mosquitoes without *Wolbachia* (11–13, 28–30).

Additionally, the proportion of *w*Mel-*Ae. aegypti* with detectable levels of DENV in the saliva is typically negligible (10, 29, 31). However, ‘breakthrough’ infections, where mosquitoes develop infectious saliva despite the presence of *Wolbachia*, occur more frequently when lab-reared mosquitoes are fed viremic blood from dengue patients (11–13, 32). Various studies have also demonstrated that *w*Mel is susceptible to heat in *Ae. aegypti*, particularly at the larval stage. Specifically, when *w*Mel-*Ae*. *aegypti* are reared at an average daily temperature above 30°C *w*Mel density is reduced (33–35) and the susceptibility of *w*Mel-*Ae. aegypti* to DENV serotype 2 infection increases in the laboratory (36).

Here we explore this relationship further by using viremic blood from dengue patients to examine the effect of a high temperature rearing regime on breakthrough infections in *w*Mel-*Ae. aegypti*. We reared WT and *w*Mel-*Ae. aegypti* at standard (28 ± 4°C) or elevated (31 ± 4°C) temperatures, fed them viremic blood from patients with confirmed DENV-1, -2, or 4 infections, then assessed the impact of rearing temperature on breakthrough infections. We also assessed whether DENV RNA and *w*Mel density in the mosquito head/thorax tissues were associated with the risk of mosquitoes developing infectious saliva.

## Methods

### Patient enrolment and diagnostics

All study participants described here were virologically-confirmed dengue inpatients at the Hospital for Tropical Diseases (HTD) in Ho Chi Minh City, Viet Nam. The protocol enabling participant enrolment was approved by the Scientific and Ethical Committee of the HTD and University of Oxford Tropical Research Ethical Committee (approval numbers: HTD EC CS/NÐ/14/12 and OxTREC 45-14). The protocol allowed for enrolment and collection of venous blood from patients admitted to the HTD who were suspected to have dengue, were >15 years of age and with <96 hours of signs or symptoms. Written informed consent was obtained from all patients participating in the study and was taken by qualified staff from the HTD. The exclusion criteria were unconscious/severely ill or pregnant women.

For all enrolled patients, venous blood was drawn into an EDTA tube. An aliquot of the blood was taken to test for DENV infection by NS1 rapid test and, if positive, then tested using PCR to determine the serotype and estimate the DENV RNA concentration (RNAemia concentration) (12). RNAemia concentration is reported here as genome copies per mL, where the ratio between genome copies per mL and plaque forming units per mL was previously determined as 214:1 for DENV-1, 73:1 for DENV-2, and 101:1 for DENV-4 (37). Thirteen virologically confirmed DENV cases were enrolled including 3 x DENV-1, 8 x DENV-2 and 2 x DENV-4.

### Mosquito rearing

The wild type (WT) *Ae. aegypti* laboratory colony derived from field collected mosquitoes in Ho Chi Minh City (HCMC), Viet Nam, has been previously described (38). The *w*Mel-infected line was generated after backcrossing the WT males from HCMC with *Wolbachia*-infected females from Cairns (Australia) for five generations. Once the backcrossing was complete, every second generation of both colonies were outcrossed with 10% of fresh field-collected males to maintain the genetic background similar to the local field population.

Colony maintenance was performed at ∼28°C, 75–85% relative humidity (RH) and 12h:12h light-dark cycle with access to 10% sucrose solution *ad libitum*. For experiments, 300 mosquito larvae were placed in plastic trays (17 × 12 × 7 cm) with 500mL of tap water and food. The larvae were reared on a limited amount of food, with a regime of 25 mg of fish food (Thức Ăn Cá, KaoKui, Viet Nam) daily, equivalent to 10% of the food given for the colony maintenance. Pupae were collected daily and retained in plastic cups in each temperature regime. After emergence, mosquitoes had access to water and 10% of sucrose solution *ad libitum*.

### Rearing temperature regimes

To assess the impact of elevated temperature on DENV transmission by *w*Mel-*Ae. aegypti* we adopted a rearing temperature profile that exceeds the severe daily mean temperature threshold for Cairns, Australia. This is defined as 30.4°C based on temperature data from 1958-2011 (39). WT and *w*Mel-*Ae. aegypti* were reared at a mean temperature of 31°C with a Diurnal Temperature Range (DTR) of 8°C (31 ± 4°C). Controls were reared at a mean temperature of 28°C with a DTR of 8°C (28 ± 4°C). Fluctuating temperature regimes followed a sinusoidal progression during the day and a negative exponential decrease at night. The maximum and minimum temperatures were reached at 12:00 and 00:00, respectively. The mosquitoes from both lines were hatched and placed in two different incubators (Percival Scientific Inc, USA) that maintained the respective temperature regimes at ∼70-80% RH, with 12:12 hours light: dark cycle. The fluorescent lights were scheduled to turn on at 06:00 and off at 18:00. Temperature regimes were maintained through all mosquito life stages including after viremic blood feeds. Temperature and humidity data loggers (Lascar Electronics, EL-USB-2, UK) recorded temperatures hourly in two locations in the incubators (top and bottom) to ensure there were minimal differences in temperature according to their locations. Twice a day between 9am and 5pm, additional thermometers (Up Aquarium Supply, China) were placed in 2 trays of each mosquito line containing the larvae to compare water temperature to the air temperature displayed in the incubators. Actual air temperatures within the incubators matched the programmed temperature profile. The extracted data of each day during all the experiments were plotted together and minimal variation was observed (Supp. Fig. 1).

### Mosquito blood feeding and harvesting

Female mosquitoes (3-5-days old) from both rearing conditions were starved for 24 h prior to viremic blood challenge. WT and *w*Mel-infected *Ae. aegypti* females were exposed to viremic blood in parallel for 30 minutes using artificial membrane feeders. Mosquitoes were then anesthetised on ice. All fully engorged females from each group were sorted and up to five of them were placed in small plastic cups with 5 males and covered with mesh. Water and 10% sucrose solution were provided *ad libitum* until harvesting on day 14. A maximum of 20 females per group were processed. If more than 20 females were present at day 14, then 20 females from each group were randomly selected and the remaining ones were discarded. To test for transmission potential, saliva was collected from each individual mosquito for inoculation into five naïve WT *Ae. aegypti* as described previously (11). The inoculated mosquitoes were maintained at 28 °C, 70–80% RH, and a 12:12 h light:dark cycle, with access to sucrose solution ad libitum. Each group of 5 injected mosquitoes were then harvested as a single pool 7 days post infection and processed to determine a binary positive/negative score for infectious DENV in the saliva from the donor mosquito. Note that saliva-injected mosquitoes were only tested if the head/thorax of the donor mosquito was determined to be DENV-positive.

Wing measurements of blood fed females were made after saliva was collected and was performed by two independent technicians. Mosquito dissections were performed after saliva collection by separating the head/thorax from the abdomen. Head/thorax tissues were stored individually at −80°C until processed for DENV and *Wolbachia* density quantification.

### PCR Processing

For non-blood fed mosquitoes reared at different temperatures, *Wolbachia* density was measured in 24 female mosquitoes collected at 3-5 days post-emergence. This experiment was repeated in three independent replicates, for a total of 72 mosquitoes. Females were dissected to separate the head/thorax and the abdomen. For abdomen samples, ovaries were removed and assessed separately since this tissue is heavily enriched for *w*Mel (29) and may mask differences in *Wolbachia* density in other abdomen tissues. Head/thorax, abdomen and ovaries were individually processed.

All harvested mosquitoes, dissected head/thorax or abdomen tissues, as well as saliva-inoculated naïve mosquitoes, were homogenized in a 96-well plate containing 100 μL of squash buffer (10mM Tris base, 1mM EDTA, 50mM NaCl [pH 8.2], and 0.4 mg/ml proteinase K), using 2.3 mm diameter glass beads in a beadbeater (TissueLyser II, Qiagen), then heated at 56°C for 10 min and 98°C for 15 min, as previous described (12).

Samples were clarified by centrifugation then 2 μL of sample was added to two separate PCR reaction mixes: the first reaction was a RT-qPCR performed to quantify tissue-specific DENV levels using pan-serotype primers (37), relative to an *Ae*. *aegypti* reference gene, *RPS17.* The second reaction was a qPCR used to quantify *w*Mel per cell (*w*Mel density), where the *WSP* gene of *Wolbachia,* and *RPS17* were used, as previously described (40, 41). Both relative DENV levels and *Wolbachia* density were calculated using the delta Ct method (2^CT(reference)^/ 2^CT(target)^). Across both PCRs, the internal *RPS17* control had to be positive to accept the results as valid. For virus transmission experiments, if the head/thorax tissue of a female was positive for DENV at day 14 post blood feeding, the five saliva-injected naïve mosquitoes were harvested at 7 days post infection, pooled and also subjected to RT-qPCR. If the head/thorax tissue of a female was negative, all inoculated naïve mosquitoes were discarded.

### Data analyses

Marginal logistic regression models were used to determine the odds of mosquitoes developing an infection in their head/thorax, and for developing infectious saliva. The model was adjusted for the additional effects of rearing temperature, virus serotype and RNAemia titre (included in the model as a log_10_ transformation) on the DENV infection status. An interaction term between temperature and mosquito type (WT or *w*Mel) was included in the model.

Values for DENV levels and *w*Mel density in mosquito tissues were not normally distributed, and values for DENV levels were also zero-inflated. Kruskal-Wallis test with Dunn’s correction was used to compare the relative DENV levels in the head/thorax of WT and *w*Mel *Ae. aegypti* reared at standard or elevated temperatures. Mann-Whitney U tests were used to compare *w*Mel densities in *Ae. aegypti* tissues.

For determining the odds of *w*Mel-*Ae. aegypti* developing infectious saliva, a marginal logistic regression model was adjusted for the effects of rearing temperature, virus serotype, DENV RNA levels in the head/thorax, *w*Mel density in the head/thorax, and RNAemia titre (included in the model as a log_10_ transformation).

Analyses for the adjusted marginal logistic regression models were performed with the statistical software R, version 4.3.3 (R Foundation for Statistical Computing).

## Results

### Elevated rearing temperature reduces wMel’s protection against patient-derived DENV serotypes 1, 2 and 4

We examined the effect of temperature on *w*Mel-mediated inhibition of DENV transmission in *Ae. aegypti*, using blood from patients with virologically confirmed DENV-1, DENV-2, or DENV-4 infection (Supp. Table 1). At 14 days post-infection, nearly all wild-type (WT) mosquitoes developed DENV infection in the head/thorax (a proxy for infection and dissemination), irrespective of rearing temperature (98.0% at 28°C ± 4°C and 95.6% at 31°C ± 4°C; Table 1).

**Table 1.**
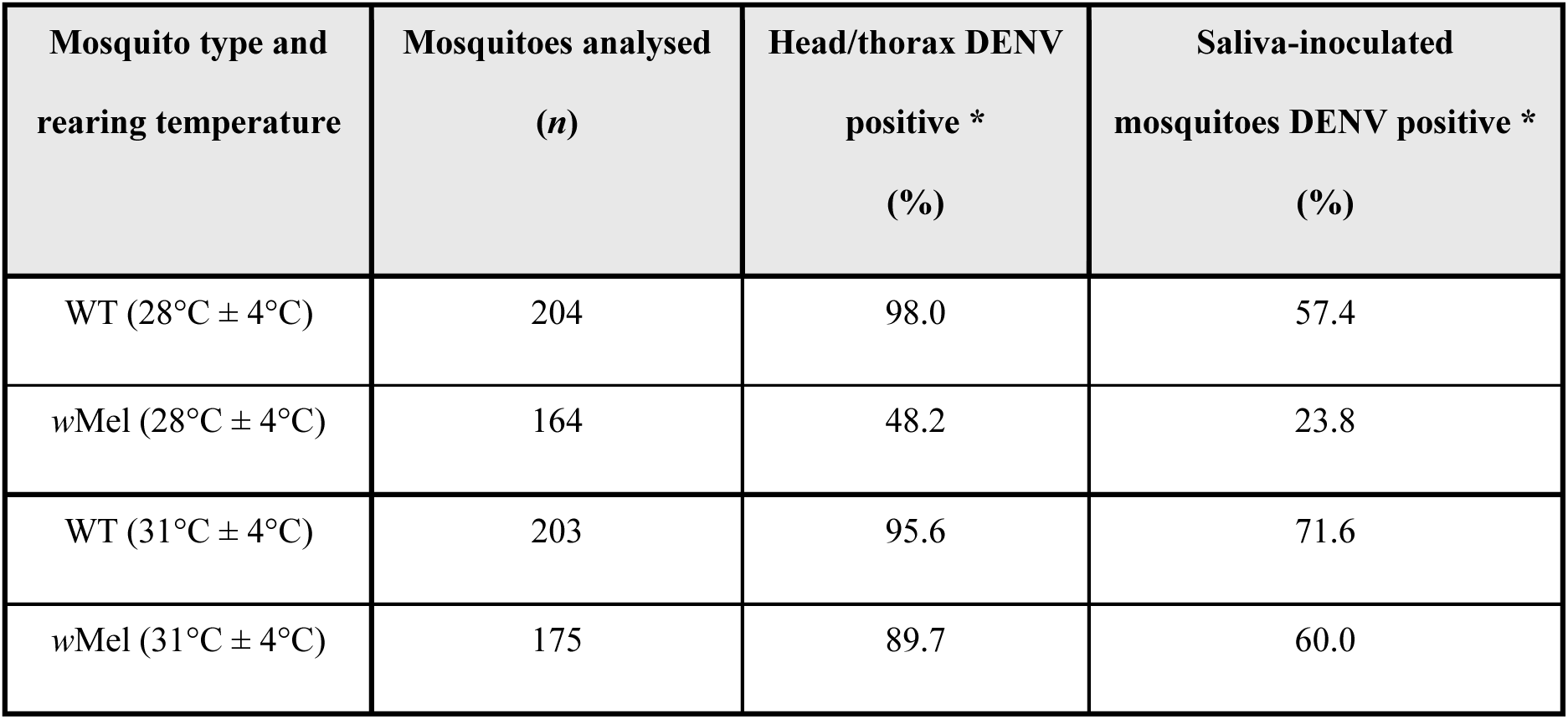
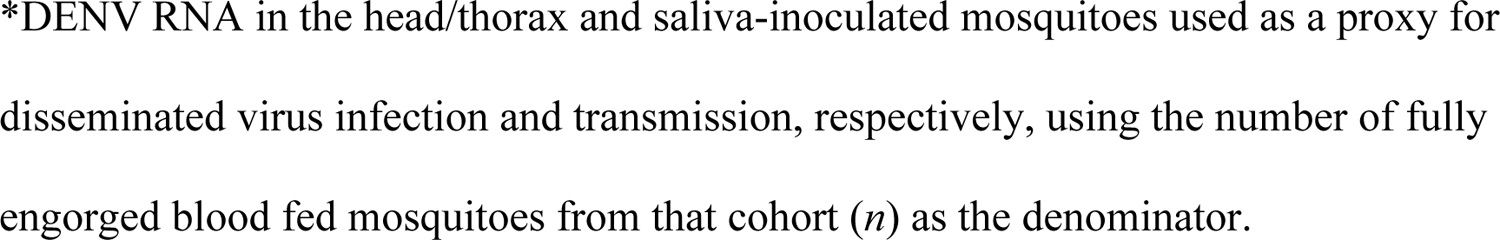
DENV infection and dissemination in, and transmission by, WT and *w*Mel-*Ae. aegypti* fed on viremic blood.

| Mosquito type and rearing temperature | Mosquitoes analysed (*n*) | Head/thorax DENV positive * (%) | Saliva-inoculated mosquitoes DENV positive * (%) |
| --- | --- | --- | --- |
| WT ($28^{\circ}\text{C} \pm 4^{\circ}\text{C}$) | 204 | 98.0 | 57.4 |
| *w*Mel ($28^{\circ}\text{C} \pm 4^{\circ}\text{C}$) | 164 | 48.2 | 23.8 |
| WT ($31^{\circ}\text{C} \pm 4^{\circ}\text{C}$) | 203 | 95.6 | 71.6 |
| *w*Mel ($31^{\circ}\text{C} \pm 4^{\circ}\text{C}$) | 175 | 89.7 | 60.0 |

\*DENV RNA in the head/thorax and saliva-inoculated mosquitoes used as a proxy for disseminated virus infection and transmission, respectively, using the number of fully engorged blood fed mosquitoes from that cohort ( $n$ ) as the denominator.

In contrast, disseminated infection rates in *w*Mel-infected *Ae. aegypti* were substantially lower when mosquitoes were reared at 28°C ± 4°C (48.2%), but increased markedly when reared at 31°C ± 4°C (89.7%). The risk of DENV infection in the head/thorax was determined using a marginal logistic regression model accounting for mosquitoes’ *w*Mel infection status, rearing temperature, bloodmeal DENV serotype, and bloodmeal RNAemia titre. A significant interaction was detected between *w*Mel infection status and rearing temperature (*p* < 0.001).

*w*Mel infection was associated with a large reduction in the odds of disseminated DENV infection for mosquitoes reared at 28°C ± 4°C (adjusted odds ratio [aOR] = 0.01, 95% CI: 0.00–0.03, *p* < 0.001) and also for mosquitoes reared at 31°C ± 4°C ( aOR = 0.37, 95% CI: 0.15–0.88, *p* = 0.024; Fig. 1A, Supp. Table 2).

**Fig. 1.**
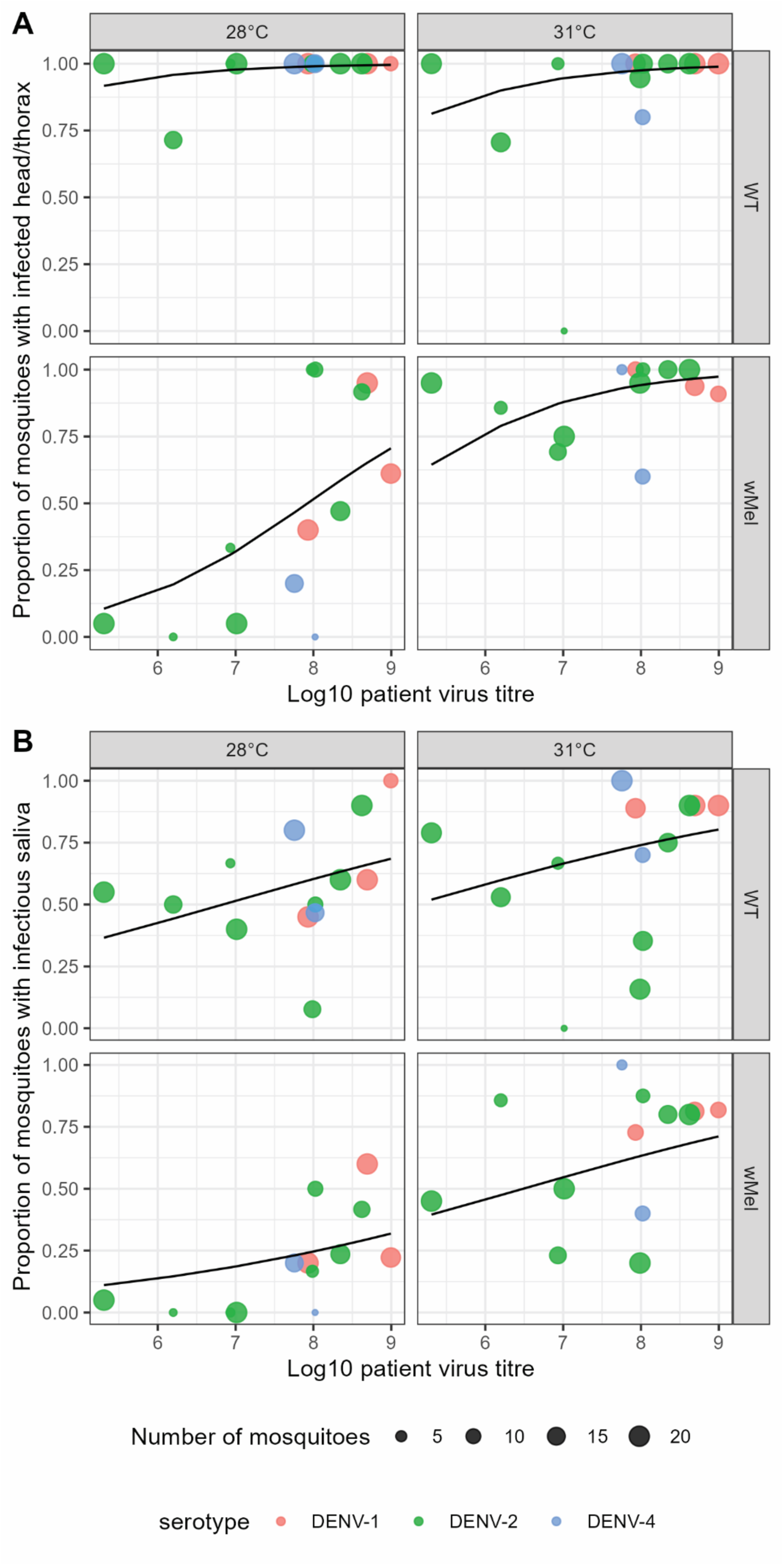
Proportion of WT and *w*Mel-*Ae. aegypti* with viral RNA in the head/thorax or infectious saliva when reared at standard or elevated temperatures. *Ae. aegypti* infected with *w*Mel or uninfected (WT) were reared at 28 or 31°C (± 4°C) and fed indirectly on virologically confirmed DENV-1, -2, or -4 patient blood. Dots represent the proportion of each cohort positive for DENV in the head/thorax (A), or in naïve mosquitoes that were inoculated with saliva collected from index mosquitoes (B). Data are plotted as a function of log_10_ plasma RNAemia (DENV RNA copies per milliliter) with logistic curves overlaid. The size of the dots represents the number of mosquitoes tested in each cohort (maximum of 20) and the colour indicates the viremic blood DENV serotype.

**Table 2.** Adjusted marginal logistic regression model for the risk of *w*Mel-*Ae. Aegypti* developing infectious saliva.

| Characteristic | Adjusted Odds Ratio | 95% CI | *p*-value |
| --- | --- | --- | --- |
| DENV RNA level<sup>l</sup> | 4.37 | 2.09 – 10.6 | <0.001 |
| *w*Mel density<sup>l</sup> | 0.90 | 0.56 – 1.42 | 0.7 |
| Temperature |  |  |  |
| 28°C ± 4°C | Ref | - | - |
| 31°C ± 4°C | 1.62 | 0.80 – 3.29 | 0.2 |
| Log<sub>10</sub>(patient RNAemia)<sup>l</sup> | 0.83 | 0.56 – 1.21 | 0.3 |
| Serotype |  |  |  |
| DENV-1 | Ref | - | - |
| DENV-2 | 0.35 | 0.15 – 0.76 | 0.009 |
| DENV-4 | 2.34 | 0.52 – 16.8 | 0.3 |

<sup>l</sup>aOR represents an increase in odds associated with a one unit increase in each characteristic. For DENV RNA levels and *w*Mel density these are derived from mosquito head/thorax tissues.

Rearing temperature did not significantly affect the odds of disseminated infection in WT mosquitoes (reference: 28°C ± 4°C; *p* = 0.126). However, *w*Mel-infected mosquitoes reared at 31°C ± 4°C had significantly higher odds of developing a disseminated infection compared with those reared at 28°C ± 4°C (aOR = 16.57, 95% CI: 8.32–32.99, *p* < 0.001).

As expected, when reared at 28 °C ± 4 °C, a smaller proportion of *w*Mel–*Ae. aegypti* developed a breakthrough (BT) infection—defined as the presence of infectious saliva—compared with WT mosquitoes (23.8% vs 57.4%; Table 1). When reared at 31 °C ± 4 °C, the proportion of *w*Mel–*Ae. aegypti* with BT infection increased, but remained lower than that observed in WT mosquitoes (60.0% vs 71.6%).

These findings were supported by a marginal logistic regression model accounting for rearing temperature, virus serotype, and RNAemia titre. Consistent with the head/thorax DENV infection model, there was a significant interaction between *w*Mel infection status and rearing temperature (*p* < 0.001). Strong inhibition of BT infection was observed in *w*Mel–*Ae. aegypti* relative to WT mosquitoes when reared at 28 °C ± 4 °C (aOR = 0.20, 95% CI = 0.13–0.32, *p* < 0.001; Fig. 1B). An inhibitory effect of *w*Mel on BT infection was also observed in mosquitoes reared at 31 °C ± 4 °C, but was attenuated (aOR = 0.64, 95% CI = 0.41–1.00, *p* = 0.05).

A higher rearing temperature significantly increased the odds of BT infection for both WT mosquitoes (31 °C ± 4 °C vs 28 °C ± 4 °C aOR = 1.89, 95% CI = 1.24–2.89, *p* = 0.003) and *w*Mel–*Ae. aegypti* (aOR = 6.04, 95% CI = 3.68–9.90, *p* < 0.001).

Virus serotype also influenced BT infection risk. Compared with mosquitoes fed DENV-1, those fed DENV-2 had significantly lower odds of developing infectious saliva (aOR = 0.49, 95% CI = 0.32–0.75, *p* = 0.001), whereas no difference was observed for DENV-4 (*p* = 0.638). In addition, and consistent with previous reports, higher DENV RNAemia titres were associated with increased odds of BT infection (aOR per log_10_ increase in RNAemia = 1.25, 95% CI = 1.06–1.48, *p* = 0.008) (11).

Having established that rearing mosquitoes at 31°C (± 4°C) increases the odds of disseminated and BT infections occurring in *w*Mel-*Ae. aegypti*, we next asked whether temperature also influenced the amount of DENV RNA in the head/thorax in *w*Mel-*Ae. aegypti* versus WT mosquitoes. Compared to WT, *w*Mel-*Ae. aegypti* had significantly less DENV in the head/thorax when reared at 28°C (± 4°C) (Fig. 2A, B, C; *p* < 0.001, Kruskal-Wallis with Dunn’s correction). However, the magnitude of *w*Mel’s protection was reduced in mosquitoes reared at 31°C (± 4°C), and only significantly reduced DENV-2 levels compared to WT. Notably, the amount of DENV in the head/thorax was significantly higher in *w*Mel-*Ae. aegypti* reared at 31°C (± 4°C) compared to 28°C (± 4°C) (*p* < 0.0001 for DENV-1 and -2, *p* <0.01 for DENV-4).

**Fig. 2.**
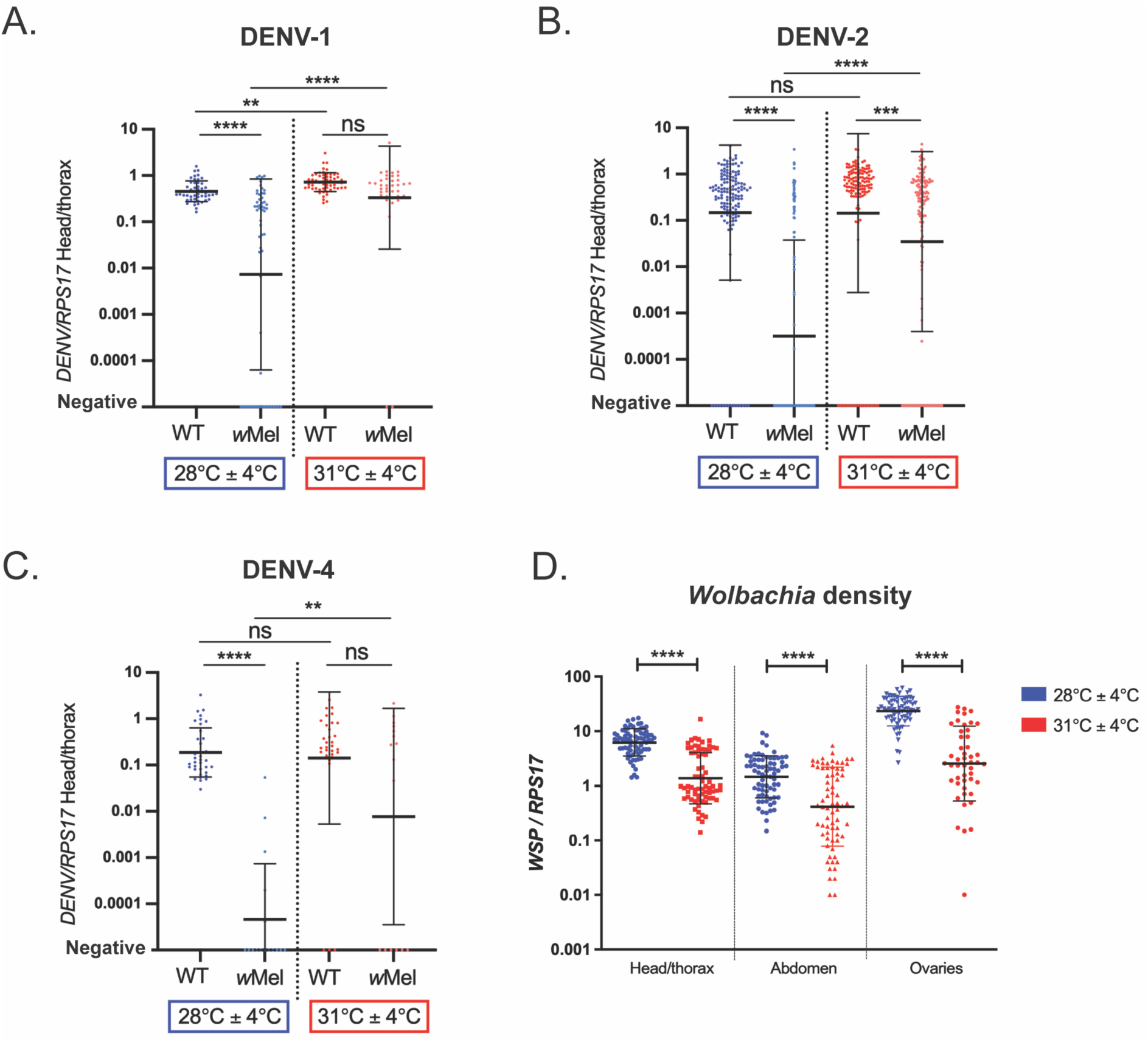
Elevated temperatures reduce the magnitude of *w*Mel’s protection against DENV-1, -2 and -4, and reduce *w*Mel density. (A, B, C) 3–5-day old female WT and *w*Mel-*Ae. aegypti* were fed blood from virologically confirmed dengue patients. Fully engorged females were selected and then incubated for 14 days. Each mosquito was dissected to separate the head/thorax which was then assessed for DENV quantification by RT-qPCR. Data are derived from Supp. Table 1, where each data point is from a single blood fed, dissected mosquito. The geometric mean and geometric standard deviation are shown. Statistical analyses were performed using Kruskal-Wallis test with Dunn’s correction for multiple comparisons, where \*\**p* < 0.01, \*\*\**p* < 0.001, \*\*\*\**p* < 0.0001 and *ns* indicates not significant. **(D)** Female mosquitoes were hatched and reared at 28°C (± 4°C) or 31°C (± 4°C), then collected 3-5 days post-emergence. Up to 72 female mosquitoes were dissected across 3 independent experiments, and the abdomen, head/thorax and ovaries were individually assessed for the number of *w*Mel per cell by qPCR. Each data point is from a dissected individual mosquito. The geometric mean and geometric standard deviation are shown. Statistical analyses were performed using Mann-Whitney test, where \*\*\*\**p* < 0.0001.

It has previously been reported that elevated rearing temperatures can reduce the density of *w*Mel in *Ae. aegypti* (34, 36), and we next confirmed that the elevated rearing temperature regime used here was sufficient to reduce the amount of *w*Mel in the head/thorax, abdomen, and ovaries of mosquitoes. Mosquitoes reared at 31°C (± 4°C) were found to have approximately 1log_10_ less *w*Mel than those reared at 28°C (± 4°C) in each of the tissues examined (Fig. 2D; *p* < 0.0001, Mann-Whitney test). We also measured significantly reduced wing length in WT and *w*Mel-*Ae. aegypti* reared at 31°C (± 4°C) compared to their counterparts reared at 28°C (± 4°C) (Supp. Fig. 2; *p* < 0.0001, Kruskal-Wallis test with Dunn’s correction) but there was no difference in wing length of *Ae. aegypti* with or without *w*Mel when reared at the same temperature (*p* > 0.999).

### Breakthrough infections in wMel-Ae. aegypti reared at standard or elevated temperatures can be predicted by DENV RNA levels, but not wMel density, in head/thorax tissues

We next considered whether the difference between *w*Mel-*Ae. aegypti* that developed infectious saliva and those that did not could be explained by the amount of DENV RNA or *w*Mel density in the head/thorax tissues of these mosquitoes. We tested this using a marginal logistic regression model adjusted for DENV serotype, DENV RNA levels in the head/thorax, *w*Mel density in the head/thorax, temperature, and patient RNAemia. For every one unit increase in DENV RNA levels in the head/thorax, the odds of *w*Mel-*Ae. aegypti* developing infectious saliva increased by 4.37 times (95% CI = 2.09 - 10.6, *p* <0.001) (Table 2). The model did not find *w*Mel density, temperature or patient RNAemia to have a significant association with BT infection. This indicates that of the characteristics tested, the amount of DENV that establishes in the mosquito head/thorax may be the most important determinant in whether *w*Mel-*Ae. aegypti* develop infectious saliva.

## Discussion

Whilst a substantial body of evidence supports the effectiveness of *Wolbachia* introgression for biocontrol of dengue, there remains a risk this intervention may fail in some settings.

*Wolbachia* strain *w*Mel has previously been reported to be susceptible to high temperatures, presenting a potential climatic or short-term limitation to its effectiveness (34, 42). First instar larvae are thought to be the most vulnerable life stage to temperature-induced loss of *w*Mel, so if *Ae. aegypti* eggs are laid in sun-exposed water there is a risk that adults may emerge without *Wolbachia* (33, 42). Additionally, females that emerge with low or no *w*Mel in their ovaries may experience breakdown of maternal transmission of the bacterium, potentially causing *Wolbachia* to drop out of populations. Work from Ross *et al*., suggests heatwaves may only transiently reduce the frequency of *w*Mel in *Ae. aegypti* populations where the bacterium has been stably established for prolonged periods (33). However, in regions where *w*Mel is being introduced into populations for the first time, or only recently introduced, such as in Nha Trang City, Viet Nam, high ambient temperatures may interfere with sustained introgression leading to spatial heterogeneity and ultimately hampering efforts to control DENV using this approach (43).

As shown here, and in prior studies, in addition to affecting the prevalence of *w*Mel-infected individuals within *Ae. aegypti* populations, elevated rearing temperatures can also reduce the density of *w*Mel in *Ae. aegypti* tissues. Mancini *et al.,* showed that heat treatment of *w*Mel-*Ae. aegypti* (reared with a diurnal rearing temperature with a minimum of 27°C, maximum of 37°C) during the larval, adult, or throughout all life stages significantly reduced the density of *w*Mel in the midgut and salivary glands - critical anatomical tissues for DENV infection and transmission, respectively (36). Interestingly, the authors found that when this heat treatment was applied at the larval stage only, *w*Mel density significantly, albeit incompletely, recovered by 14 days post emergence. Here, we applied an equivalent diurnal rearing temperature range across all mosquito life stages and similarly reported a significant drop in the density of *w*Mel in head/thorax, abdomen (ovary-extracted) and ovary only tissues. Mancini *et al.,* used a cell culture grown DENV-2 strain (New Guinea C) to show that heat treatment led to a significant increase in the number of *w*Mel-*Ae. aegypti* that acquired a DENV-2 infection compared to a matched control cohort. Additionally, their results indicated that heat treated *w*Mel-*Ae. aegypti* no longer protect against DENV-2 transmission (36).

We extend this prior work by assessing the effects of elevated rearing temperature on the infection and transmission potential of *w*Mel-*Ae. aegypti* fed patient blood from viremic DENV-1, -2 and -4 patients. Using this natural history model, our data corroborates that a higher rearing temperature (27°C – 35°C, compared to 24°C - 32°C) increases the risk of *w*Mel-*Ae. aegypti* acquiring a disseminated infection and developing infectious saliva. Our results also confirm previous findings that elevated rearing temperature significantly increases the probability of *Wolbachia*-uninfected *Ae. aegypti* developing infectious saliva (44–46). However, in this study, *w*Mel still significantly reduced the risk of *w*Mel-*Ae. aegypti* developing infectious saliva compared to matched WT mosquitoes, even at the higher rearing temperature.

Our data showed that DENV RNA levels were significantly increased in the head/thorax of heat-treated *w*Mel-*Ae. aegypti,* and that the probability of *w*Mel-*Ae. aegypti* mosquitoes developing infectious saliva was significantly associated with higher levels of DENV RNA, but not lower *w*Mel density, in these tissues. This may suggest that breakthrough infections occur more frequently in *w*Mel-*Ae. aegypti* reared at higher temperatures due to elevated levels of DENV replication. It is also possible that temperature causes critical differences in *w*Mel density to occur in defined regions of the head/thorax such as the salivary glands which were not specifically examined in this study.

The increased risk of DENV transmission under elevated temperatures is important to consider, as enhanced replication of DENV in the presence of *w*Mel theoretically provides an opportunity for the selection of viral variants with increased transmission potential. However, in order for DENV to emerge as resistant to the intervention, these variants would also need to remain fit in humans. Given the substantial evolutionary bottlenecks reported for DENV that occur at both the vertebrate and invertebrate infection stages, there are still likely to be significant barriers to resistant viruses emerging (25).

Here, we considered a scenario where *w*Mel-*Ae. aegypti* were exposed to high temperatures everyday across a three-week period, with no opportunity for recovery of *w*Mel density at lower temperatures. In reality, if such a heat wave were to occur in the field, adult mosquitoes would likely seek to rest in shady areas with the lowest available temperature (47) perhaps facilitating some recovery of *w*Mel density and reducing the risk of DENV transmission.

Regardless, it is important to understand the limitations of using *w*Mel in geographic regions that experience prolonged heat events, and our results suggest that we can expect to see more instances of DENV transmission in *w*Mel-established regions when mosquitoes are consistently exposed to consecutive days with an average daily temperature above 30°C. Field release sites with established populations of *w*Mel-*Ae. aegypti* should consider implementing surveillance for the weeks immediately following such climatic events, to monitor for the increased occurrence of DENV infection in mosquitoes, and/or a loss of *w*Mel infection frequency in the mosquito population.

Additionally, in countries aiming to establish *Wolbachia* as a biocontrol method that consistently experience average daily temperatures above 30°C, it may be prudent to instead consider introgression of another antiviral *Wolbachia* strain, *w*AlbB, which has repeatedly been shown to be more tolerant of high temperatures (34, 36). Whilst *w*AlbB brings other fitness costs, such as substantially reduced fertility in females hatched following egg quiescence (48), the climate in certain locations may justify its use.

## Acknowledgements

We thank the patients and their families for their participation in this study, and the staff from the HTD. We thank Aimee Altermatt (Modelling and Biostatistics group, Burnet Institute) for statistical consultation and data visualisation support. This work was supported by the Wellcome Trust (UK) (Award No.102591/Z/13/Z), the National Health and Medical Research Council (Australia) (APP1182432) and the Australian Research Council (DP220102997).

---

**Supp. Table 1.** DENV infection (head/thorax positive) and transmission (saliva positive) in WT and *w*Mel-*Ae. aegypti* reared under standard or elevated temperatures –.

| Serotype | Patient RNA-emia titre | Head/thorax DENV positive (n) |  |  |  | Saliva DENV positive (n) |  |  |  |
| --- | --- | --- | --- | --- | --- | --- | --- | --- | --- |
|  |  | WT |  | *w*Mel |  | WT |  | *w*Mel |  |
|  |  | 28°C<br>(± 4°C) | 31°C<br>(± 4°C) | 28°C<br>(± 4°C) | 31°C<br>(± 4°C) | 28°C<br>(± 4°C) | 31°C<br>(± 4°C) | 28°C<br>(± 4°C) | 31°C<br>(± 4°C) |
| DENV-1 | 8.51 x 10<sup>7</sup> | 20<br>(20) | 18<br>(18) | 8 (20) | 11<br>(11) | 9 (20) | 16<br>(18) | 4 (20) | 8<br>(11) |
| DENV-1 | 4.91 x 10<sup>8</sup> | 20<br>(20) | 20<br>(20) | 19<br>(20) | 15<br>(16) | 12<br>(20) | 18<br>(20) | 12<br>(20) | 13<br>(16) |
| DENV-1 | 9.88 x 10<sup>8</sup> | 9 (9) | 20<br>(20) | 11<br>(18) | 10<br>(11) | 9 (9) | 18<br>(20) | 4 (18) | 9<br>(11) |
| **DENV-1 combined %** |  | **100** | **100** | **66** | **95** | **61** | **90** | **35** | **79** |
| DENV-2 | 2.03 x 10<sup>5</sup> | 20<br>(20) | 19<br>(19) | 1 (20) | 19<br>(20) | 11<br>(20) | 15<br>(19) | 1 (20) | 9<br>(20) |
| DENV-2 | 1.58 x 10<sup>6</sup> | 10<br>(14) | 12<br>(17) | 0 (2) | 6 (7) | 7 (14) | 9 (17) | 0 (2) | 6 (7) |
| DENV-2 | 8.57 x 10<sup>6</sup> | 3 (3) | 6 (6) | 1 (3) | 9 (13) | 2 (3) | 4 (6) | 0 (3) | 3<br>(13) |
| DENV-2 | 1.03 x 10<sup>7</sup> | 20<br>(20) | 0 (1) | 1 (20) | 15<br>(20) | 8 (20) | 0 (1) | 0 (20) | 10<br>(20) |
| DENV-2 | 9.70 x 10<sup>7</sup> | 13<br>(13) | 18<br>(19) | 6 (6) | 19<br>(20) | 1 (13) | 3 (19) | 1 (6) | 4<br>(20) |
| DENV-2 | 1.06 x<br>10<sup>8</sup> | 10<br>(10) | 17<br>(17) | 10<br>(10) | 8 (8) | 5 (10) | 6 (17) | 5 (10) | 7 (8) |
| DENV-2 | 2.22 x<br>10<sup>8</sup> | 20<br>(20) | 16<br>(16) | 8 (17) | 15<br>(15) | 12<br>(20) | 12<br>(16) | 4 (17) | 12<br>(15) |
| DENV-2 | 4.19 x<br>10<sup>8</sup> | 20<br>(20) | 20<br>(20) | 11<br>(12) | 20<br>(20) | 18<br>(20) | 18<br>(20) | 5 (12) | 16<br>(20) |
| **DENV-2 combined %** |  | **97** | **94** | **42** | **90** | **53** | **58** | **18** | **55** |
| DENV-4 | 5.69 x<br>10<sup>7</sup> | 20<br>(20) | 20<br>(20) | 3 (15) | 4 (4) | 16<br>(20) | 20<br>(20) | 3 (15) | 4 (4) |
| DENV-4 | 1.05 x<br>10<sup>8</sup> | 15<br>(15) | 8 (10) | 0 (1) | 6 (10) | 7 (15) | 7 (10) | 0 (1) | 4<br>(10) |
| **DENV-4 combined %** |  | **100** | **93** | **19** | **71** | **66** | **90** | **19** | **57** |

**Supp. Table. 2.** Adjusted marginal logistic regression models for the odds of viral infection in the head/thorax tissue.

| Head/thorax tissue | Adjusted odds ratio | 95% confidence interval | *p* value |
| --- | --- | --- | --- |
| (Intercept) | 0.12 | (0.01; 1.34) | 0.085 |
| WT (reference) |  |  |  |
| wMel 28°C | 0.01 | (0.00; 0.03) | <0.001 |
| wMel 31°C | 0.37 | (0.15; 0.88) | 0.024 |
| 28°C (reference) |  |  |  |
| 31°C WT | 0.39 | (0.12; 1.31) | 0.126 |
| 31°C wMel | 16.57 | (8.32; 32.99) | <0.001 |
| DENV-1 (reference) |  |  |  |
| DENV-2 | 1.00 | (0.48; 2.05) | 0.993 |
| DENV-4 | 0.24 | (0.09; 0.59) | 0.002 |
| Interaction term: wMel*31°C | 42.73 | (10.54; 173.25) | <0.001 |
| log<sub>10</sub> patient RNAemia | 2.43 | (1.83; 3.22) | <0.001 |

**Supp. Table. 3.** Adjusted marginal logistic regression models for the odds of viral infection in saliva-inoculated mosquitoes.

| Mosquitoes inoculated with saliva | Adjusted odds ratio | 95% confidence interval | *p* value |
| --- | --- | --- | --- |
| (Intercept) | 0.38 | (0.09; 1.64) | 0.196 |
| WT (reference) |  |  |  |
| wMel 28°C | 0.20 | (0.13; 0.32) | <0.001 |
| wMel 31°C | 0.64 | (0.41; 1.00) | 0.05 |
| 28°C (reference) |  |  |  |
| 31°C WT | 1.89 | (1.24; 2.89) | 0.003 |
| 31°C wMel | 6.04 | (3.68; 9.90) | <0.001 |
| DENV-1 (reference) |  |  |  |
| DENV-2 | 0.49 | (0.32; 0.75) | 0.001 |
| DENV-4 | 0.87 | (0.50; 1.54) | 0.638 |
| Interaction term: wMel*31°C | 3.19 | (1.67; 6.12) | <0.001 |
| log<sub>10</sub> patient RNAemia | 1.25 | (1.06; 1.48) | 0.008 |

**Supp. Fig. 1.**
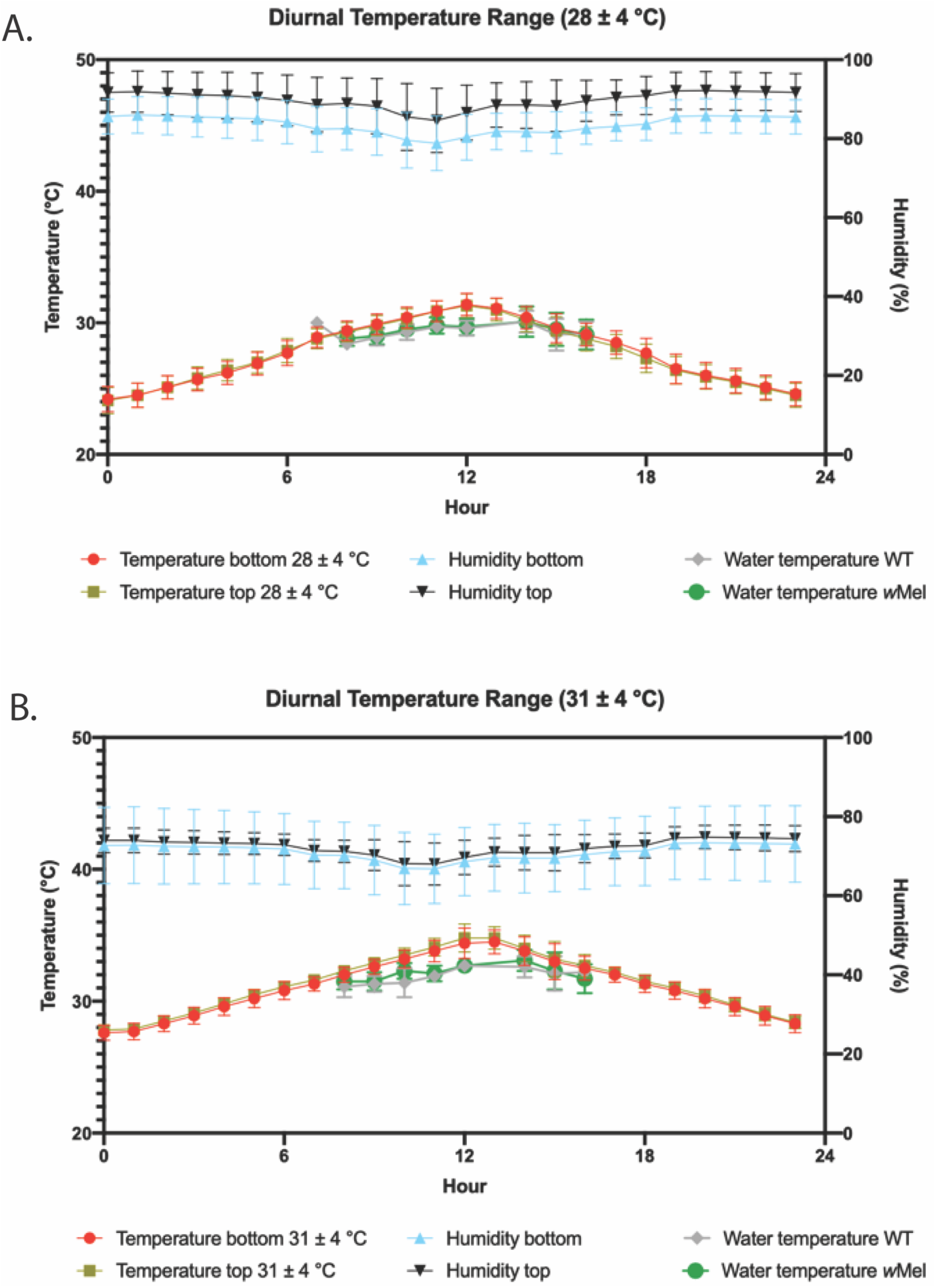
Temperature and humidity in mosquito rearing incubators. Diurnal temperature range (DTR) in the incubators is shown for **(A)** standard (28°C ± 4°C), and **(B)** elevated (31°C ± 4°C; B) rearing conditions. The lines represent the DTR at the bottom (red) and top (brown) of the incubators, as well as the humidity at the bottom (blue) and top (black). Between 9am and 5pm, the water temperature was measured for the WT (grey) and *w*Mel (green) mosquito lines. Data are the mean and standard deviation of data extracted each day during all the experiments.

**Supp. Fig. 2.**
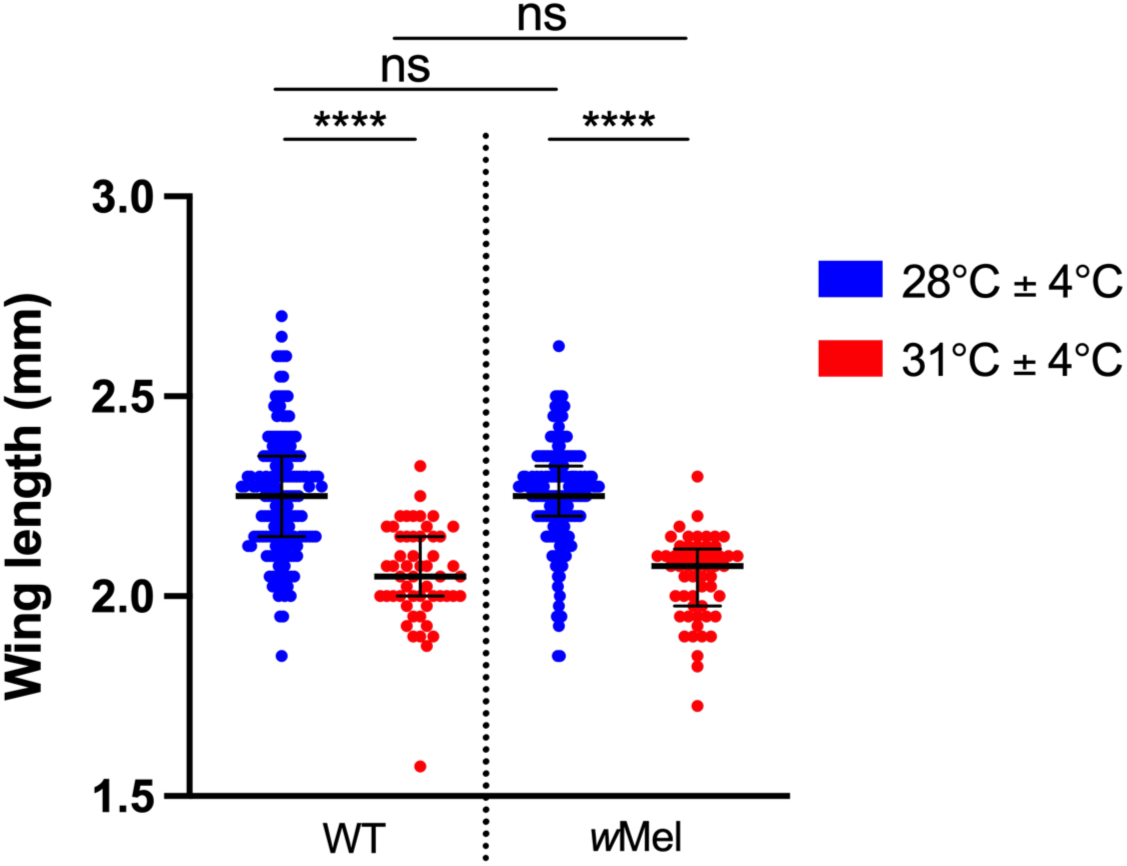
Increased rearing temperature reduces the wing length of WT and wMel-Ae. aegypti. Female mosquitoes were hatched and reared at 28°C (± 4°C) or 31°C (± 4°C), then fed on viremic blood. At 14 days post blood meal, mosquito wings were collected and measured by two independent technicians to estimate the size of each female. Data points represent individual mosquito wings and are the median and interquartile range of at least 55 wings. Statistical analysis was performed using a Kruskal-Wallis test with Dunn’s correction, where **** *p* <0.0001.

